# A Transformer-based Multi-omics Model for Translation Efficiency in *S. cerevisiae*

**DOI:** 10.64898/2026.06.21.723404

**Authors:** Dingyang Liu, Lunqiang Zhao, Yushuo Wei, Xirui Peng, Hong Lu, Xinhang Zhang, Jie Chen

**Affiliations:** College of Biomedical Engineering, Fudan University, No. 220, Handan Road, 200438, Shanghai, China; State Key Laboratory of Genetics and Development of Complex Phenotypes, Fudan University, No.220, Handan Road, 200438, Shanghai, China; Shanghai Engineering Research Center of Industrial Microorganisms, Fudan University, No. 220, Handan Road, 200438, Shanghai, China

**Keywords:** Deep Learning, Yeast Engineering, Translation Prediction, Multi-omics Model

## Abstract

Precise regulation of protein synthesis is fundamental to cellular homeostasis and remains a primary target for synthetic biology applications. However, the non-linear relationship between mRNA abundance and protein levels presents complexities that poses challenges for predictive engineering. Here, we present TRIM, a Transformer-based RNA Inference Model that leverages full-length mRNA sequences and multi-omics data to predict translation efficiency. By employing a Parallel Expert Mixer, TRIM achieves robust prediction accuracy (*R*^2^ ≥ 0.8,Pearson *r* ≥ 0.9). Trained on multimodal data from massive *Saccharomyces cerevisiae* isolates, TRIM demonstrates outstanding biological interpretability, helping to decipher complex translational patterns such as synergistic effects between bases, sequence-dependent codon preference in different stages, and distinct attention on key secondary structures. These results indicate that the integration of multi-omics data with holistic sequence modeling can effectively decode the cis-regulatory grammar of translation as well as providing a scalable and interpretable generative framework for future synthetic biology engineering.

**Availability and Implementation:** The source code and data used to produce the results and analyses presented in the manuscript are available from Github (https://github.com/ZeusLiu666/TRIM).

## 1. Introduction

Protein synthesis stands as the pivotal link between genotype and phenotype within the central dogma. Deciphering the regulatory pattern of translation is a critical prerequisite for precise protein engineering and synthetic biology applications[6,27,28]. However, transcript abundance often correlates weakly with final protein levels, suggesting that gene expression is highly modulated at the translational level[20].

Translation in eukaryotes is a highly intricate, multi-stage process governed by diverse cis-regulatory elements. It is generally divided into initiation, elongation and termination. The initiation phase is regarded as the rate-limiting step, stringently controlled by features within the 5’ Untranslated Region (5’UTR), such as secondary structures and upstream AUGs (uAUGs)[1,6,23]. During elongation, the translation rate is significantly influenced by codon usage within the Coding Sequence (CDS) and the availability of cognate tRNAs[9,15]. Finally, the 3’ Untranslated Region (3’UTR) plays a vital role in mRNA stability and avoiding degradation by recruiting trans-acting factors[17,23]. The intricate interplay among these cis-regulatory elements necessitates a holistic analysis of the full-length mRNA molecule, rather than studying isolated regions.

Deep learning has advanced significantly to dissect these complex biological networks with its capability of non-linear modeling and feature extraction. Early models like Optimus 5-Prime[19] relied on Massively Parallel Reporter Assays (MPRA). However, the synthetic libraries lack the complex endogenous genomic context and long-range interactions inherent to native genes, thus limiting their physiological relevance and generalization to natural sequences. Current translation-related models fall into two categories. The first relies on Ribosome Profiling (Ribo-seq) data, such as RiboNN[29] and STE[11]. While these models achieve high precision, their applicability is constrained by the cost and technical complexity of Ribo-seq protocols. Data scarcity limits their generalization across diverse species and tissues, especially non-model organisms with little Ribo-seq data.The second category is trained solely on mRNA sequence information, like UTR-LM[3] and UtailoR[13]. Although they reduced the reliance on expensive omics data, they often focus on isolated stages, like optimizing only the 5’UTR. This leads to performance bottlenecks due to the absence of intrinsic coupling between different stages[4,10,16]. Although some other models like mRNAdesigner[15] use full-length mRNA as input, they focuses on *de novo* sequence engineering for maximal protein expression. Consequently, their optimization often deviates from natural regulatory constraints and lacks interpretability.

To address these limitations, we introduced TRIM, a Transformer-based RNA Inference Model, that fuses proteomics and transcriptomics. TRIM utilizes readily available RNA and protein abundance data as supervision signals to predict translation efficiency (TE) directly from sequence. Crucially, our study leverages the extensive natural variation found in yeast isolates[24], marking a shift from a synthetic biology perspective to an evolutionary genetics framework. TRIM is designed to synergistically capture key motifs and structural features across 5’UTR, CDS and 3’UTR. Through such holistic modeling, TRIM delivers robust and precise predictions with remarkable biological interpretability.

## 2. Materials and Methods

### 2.1. Datasets and Preprocessing

We integrated multi-omics datasets from Teyssonnière, *et al*.[2,24], comprising paired RNA and protein abundance for 629 genes in 942 natural *Saccharomyces cerevisiae* isolates cultured in SC media (2% glucose with amino acids). We also incorporated datasets for the wild-type (WT) SUB592 (5,610 proteins) strain from Yuan Gao, *et al*.[8], and the WT BY4741 strain from Peng D., *et al*.[18].

Gene identifiers were standardized to Systematic Names using the *Saccharomyces* Genome Database (SGD) (https://www.yeastgenome.org). Full-length gene sequences were extracted based on annotations from the Eukaryotic Promoter Database (EPD) (https://epd.expasy.org/epd/). Transcriptomic data with negligible expression (TPM=0) were excluded at first. The remaining data were normalized to Transcripts per Million (TPM) and Subsequently log_2_ − transformed. For proteomics, mass spectrometry intensities were converted to Protein Expression per Million (PEM). A pseudo-count (*∈*=10^−5^) was introduced to handle zero values below the detection limit. PEM is calculated as:

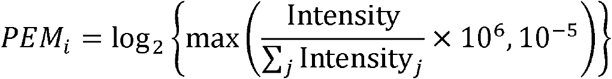

The preprocessed data were finally Z-score normalized before delivered into TRIM, and were denomalized before assessment. All of the datasets including subsets for training, validaiton, and test have already been uploaded, the download link can be found in our Github repository: https://github.com/ZeusLiu666/TRIM.

### 2.2. Input Construction

The input of TRIM consists of sequence and environmental features.

Sequences are encoded as(*L* ×5) tensors. The first four channels represent one-hot encoding of nucleotides (A,G,C,U). The fifth channel incorporates biological context: base-pairing probability for UTRs and codon bias for CDS. The pairing probability, calculated by ViennaRNA[14], refers to the cumulative pairing probability of each nucleotide with all other bases in the sequences. We mapped each triplet to a codon bias vector of the same length based on the *S. cerevisiae* codon bias table.

Environmental inputs comprise both numerical and categorical variables. Numerical features were normalized to a common scale, while categorical features were one-hot encoded, followed by a concatenation to form the environmental input vector.

### 2.3 Architecture of TRIM

The architecture of TRIM consists of the following modules (**Fig.1a**):

Feature Extraction Backbones: TRIM employs three independent Convolutional Neural Network (CNN) to respectively process the 5’UTR, CDS and 3’UTR. Each backbone consists of an initial convolution block followed by 10 Residual Blocks (kernel size = 5, output channels = 64). Variable-length sequences are processed via max pooling and subsequently aggregated into a 64-dim vector using global average pooling. Environmental features are mapped to a 64-dim vector via a Residual Multi-layer Perceptron (MLP) with a hidden dimension of 256.

**Fig. 1:**
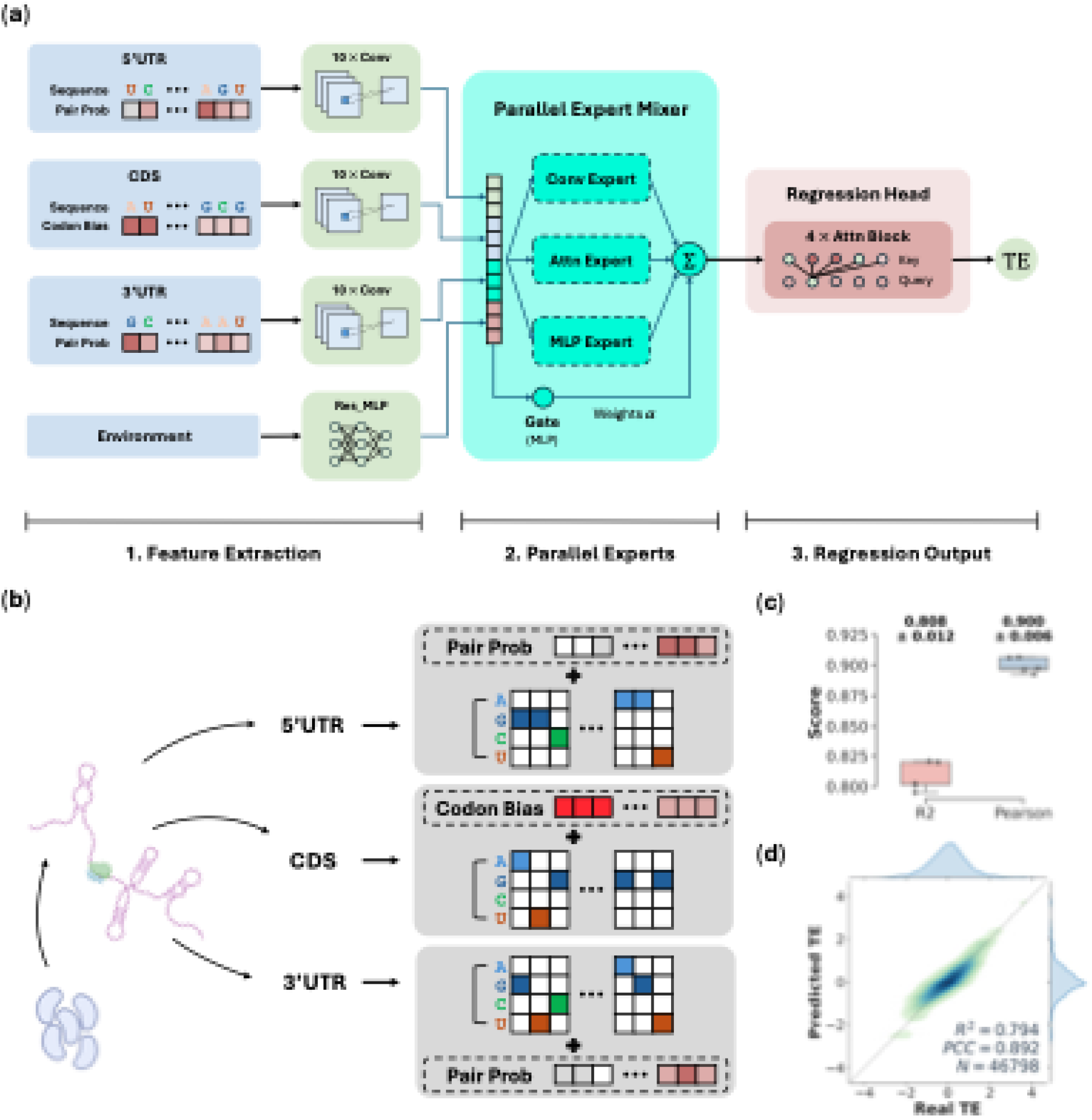
**(a)**TRIM Architecture: Feature extraction; Parallel Expert Mixer (fusing outputs from former experts via learnable gating weights *α*); Regression Head. **(b)**Multi-omic Input Integration. One-hot encoded sequences are concatenated with biological features (pairing probability and codon bias). **(c)**Five-fold Cross-validation. Dots represent data points from separate folds with the mean*±*standard deviation annotated above. **(d)**Density scatter plot of Predicted TE against Real TE. Marginal density curves represent the distribution of Real TE (top) and Predicted TE (right).

Parallel Expert Mixer: The features are concatenated and fed into the parallel expert mixer. It comprises a Convolutional Expert, an Attention Expert and an MLP Expert. The outputs of these experts (*x*_conv,_ *x*_Attn,_ *x*_MLP_) are fused using learnable weights *α*_*i*_, generated by an MLP gating network to produce the fused feature vector *F*_fused_:

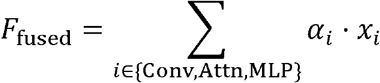

Regression Head:*F*_fused_ is put into a regression head, composed of 4 stacked transformer blocks with multi-head self-attention. The attention mechanism uses learnable matrices*W*_*q*,_*W*_*k*,_*W*_*v*,_ defining the queries, keys and values, to integrate features. The attention intensity *a*_*ij*_ of position *j*from position *i* is calculated as:

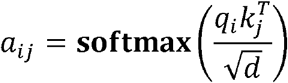

and the attention score *h*_*j*_ is further accumulated as:

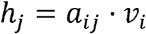

Where *q*_*i*_,*k*_*i*_,*v*_*i*_ are elements of **q**,**k**,**v**, defined as:

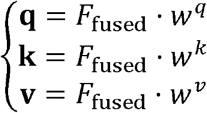

Then the scores pass through a fully connected layer to yield the final TE.

### 2.4. Hyperparameter Tunning

TRIM was implemented using a PyTorch Lightning framework and trained on an NVIDIA Tesla A100 GPU with a batch size of 768. The dataset was randomly partitioned into training (80%), validation (10%), and test (10%) sets. We employed the AdamW optimizer with a weight decay of 10^−4^. The learning rate scheduling utilized a strategy combining linear warmup with cosine annealing decay. During the cosine annealing phase, the learning rate *η*_*t*_ was defined as:

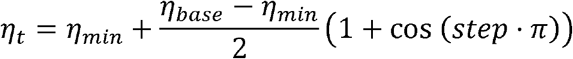

To enhance model robustness, the loss function incorporated Huber Loss with an *R*^2^-based penalty term:

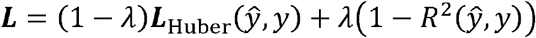

where the balancing coefficient *λ* = 0.1, and the Huber Loss parameter was set to *β* =1.5. TRIM was trained for a maximum of 100 epochs, with the best weights saved based on validation performance.

### 2.5. Sequence Optimization

We developed an *in silico* evolutionary strategy to optimize CDS for enhanced TE. We fused the CDS of reporter protein (GFP and LacZ) with high-expression UTRs (TLR061W and YLR103W) for further optimization. The process was initiated with a population of 10^6^ random synonymous variants. In each generation, the top-16 sequences with the highest TRIM-predicted TE were selected as parents and each yielded 128 offspring, each with 8 distinct synonymous substitution. The selection-mutation cycle was iterated for over 3,000 generations, after which the top-ranking sequences were harvested for analysis.

### 2.6. Regulatory Pattern Analysis

We analyzed RPL8A mutagenesis dataset from Dvir *et al*.[7], and separately grouped them based on predicted TE (top and bottom 10%). We indexed the position of each nucleotide relative to the start codon by defining the A of AUG as +1 and its upstream neighbor as −1. For every nucleotide in [−10,−1], sequences in the two subsets that harbors a specific mutation were separately grouped and their mean ΔTE from the unmutated sequence was calculated. After fixing the identified hotspot (e.g., −2C), we performed an *in silico* saturation test on secondary mutation.

## 3. Results

### 3.1. TRIM accurately predicts translation efficiency

TRIM is a deep learning model designed to accurately predict translation efficiency (TE) by jointly modeling sequence and environmental features.

During the training process, a dataset of 467,000 samples that integrates three complementary datasets was constructed to provide a comprehensive benchmark. To prevent data distortion and enhance robustness, a pseudo-count strategy was introduced to ensure numerical continuity. The resulting dataset achieves high proteome coverage as well as including strains varying from the standard laboratory strains to natural isolates, thereby mitigating biases from single platform or specific experimental conditions.

TRIM leverages a Parallel Expert Mixer layer to extract multi-omic features (**Fig.1a**). The layer consists of Convolutional Expert, Attention Expert and MLP Expert to map local, long-range and non-linear features. Since the structure of 5’UTR modulates TE by affecting the initiation rates and ribosome scanning[26], we concatenated the structural pairing probabilities and the codon bias that influences the elongation rates with raw mRNA sequences as inputs of TRIM (**Fig.1b**).

The processed dataset was randomly partitioned into training (80%), validation (10%) and test (10%) sets. Five-fold cross-validation on the test set yielded an *R*^*2*^ of 0.808 ±0.012 and a Pearson correlation coefficient (PCC) of 0.900 ± 0.006(**Fig.1c**). The density scatter plot of the validation set also demonstrated an *R*^*2*^of 0.794 with a PCC of 0.892 (**Fig.1d**), indicating that TRIM precisely captures the TE distribution of the training data. All source code can be found in our Github repository: https://github.com/ZeusLiu666/TRIM.

### 3.2. TRIM reveals *S. cerevisiae* codon bias

To evaluate the generative capability of TRIM in sequence design, we performed *in silico* evolution of CDS regions for reporter proteins (GFP and LacZ) coupled with high-expression UTRs (YLR061W and YLR103W). The optimization pipeline employed a genetic algorithm-based strategy (**Fig.2a**): starting from 10^6^ randomized synonymous variants, TRIM iteratively evolved sequences through selection-mutation cycles based on predicted TE over 3,000 generations (see Materials and Methods).

Analysis of the best-performing sequences reveals that TRIM successfully captures the intrinsic codon usage rules of *S. cerevisiae*. Although driven solely by TE maximization, the codon usage in TRIM-optimized CDS correlates highly with natural yeast codon bias (Pearson for *r* > 0.83 GFP; *r* > 0.92 for LacZ; **Fig.2bc**). Crucially, the high correlation was maintained across varying sample sizes, demonstrating TRIM’s robustness in identifying global synonymous codon preferences (**Fig.2d**).

**Fig. 2:**
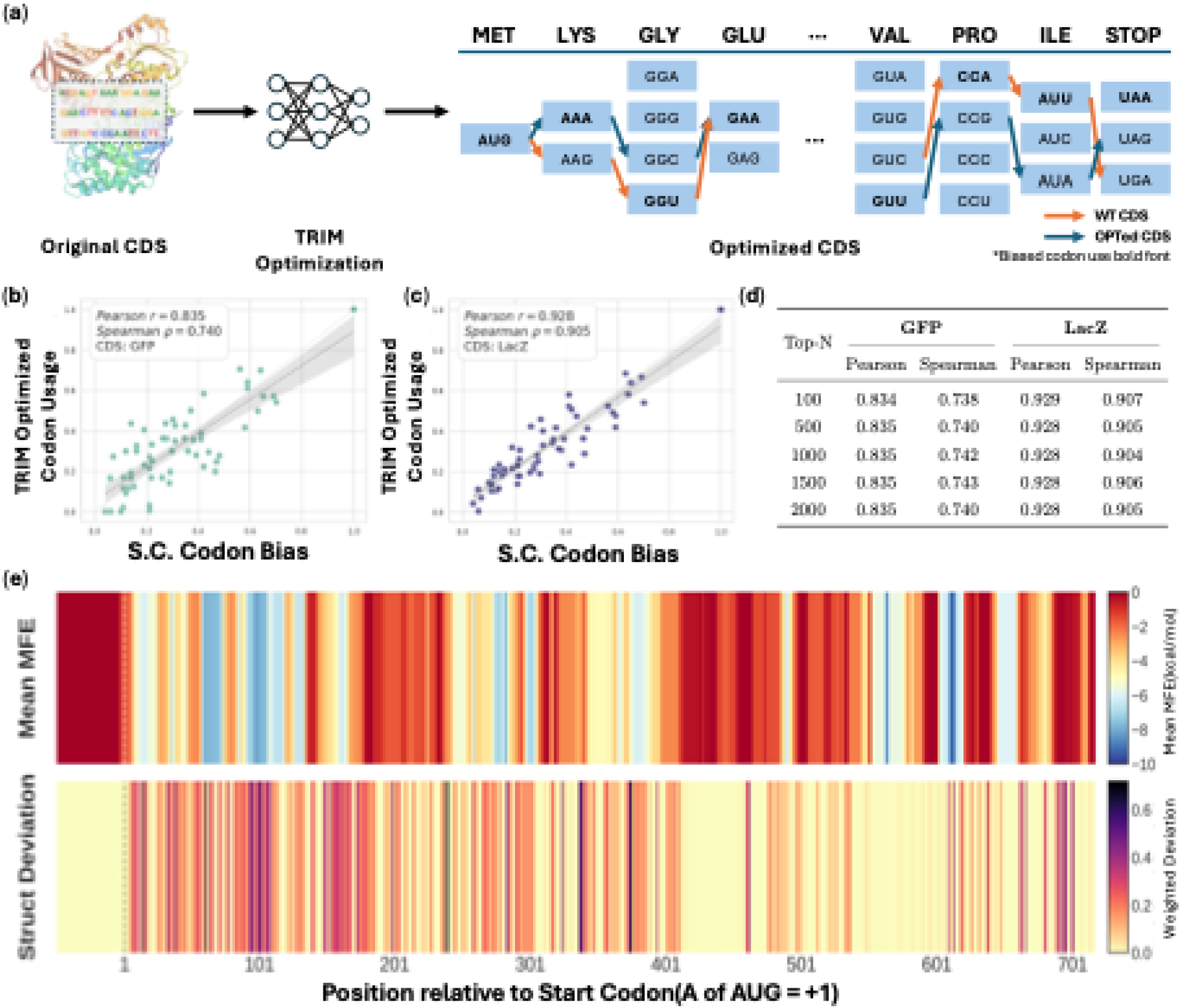
**(a)**Workflow of CDS Optimization. TRIM optimizes CDS by replacing synonymous codons. Orange arrows denote the WT codon usage, while blue arrows trace the Optimized (OPTed) codon usage. **(b, c)**Scatter plots correlating *S. cerevisiae* codon bias with TRIM-optimized codon usage frequencies in the top-2000 TE optimized CDS sequences. The gray shadow represents the 95% confidence interval. **(d)**Table of Pearson and Spearman correlation coefficients across varying numbers of top TE sequences. **(e)**Heatmap of Structural Deviation correlated with mean MFE at base-level resolution. The **x**-axis covers 50-nt 5’UTR (*x* < 0) and full-length CDS of GFP. **Upper:** MFE heatmap computed over a sliding window of 31nt. The color indicates the thermodynamic stability. **Lower:** Heatmap of quantified impact on secondary structure induced by non-preferred codon usage. The intensity is calculated as the weighted absolute difference in pairing probabilities between the employment of non-preferred codons and biased codons.

TRIM also demonstrated a deep understanding of mRNA thermodynamics. We compared the Minimum Free Energy (MFE) landscape of TRIM-optimized sequences against those composed purely of optimal codons. The optimized sequences exhibit higher mean MFE values, indicating the reduced thermodynamic stability (**Fig.2e, upper panel**). Furthermore, the analysis of codon choices revealed that non-preferred codons were not randomly distributed. Instead, they were preferentially enriched in regions that influence the formation of secondary structures. The structural deviation map (**Fig.2e, lower panel**) confirms that TRIM strategically sacrifices local codon optimality to disrupt folding elements, thereby optimizing the global landscape for efficient translation.

More notably, TRIM-optimized sequences exhibit a reduced Codon Adaptation Index (CAI) at the 5’ end of the CDS, reproducing the “Translation Ramp” observed in endogenous *S. cerevisiae* genes[21,25]. Comparative analysis showed a distinct reduction in optimal codon usage, with the overall frequency falling from 51.46% (WT) to 41.76% and a steeper reduction from 56.25% to 30.33% in the first 51nt. While the inherent cause of this ramp remains under debate, its spontaneous emergence within optimized sequence subsets suggests that TRIM has internalized global genomic constraints that extend beyond local codon optimality to modulate translation kinetics dynamically.

### 3.3. TRIM recognizes structure-sensitive translation patterns

Translation initiation is widely regarded as the rate-limiting step in eukaryotic protein synthesis. Structural motifs exert multifaceted effects on translation dynamics (**Fig.3a**): some act as physical barriers that impede ribosomal scanning, while others function as docking sites to recruit RNA-Binding Proteins(RBPs) that either enhance translation or stabilize the transcript against degradation[12]. We analyzed TRIM’s sensitivity to crucial structural regions and biophysical features from both thermodynamic and structural perspectives.

**Fig. 3:**
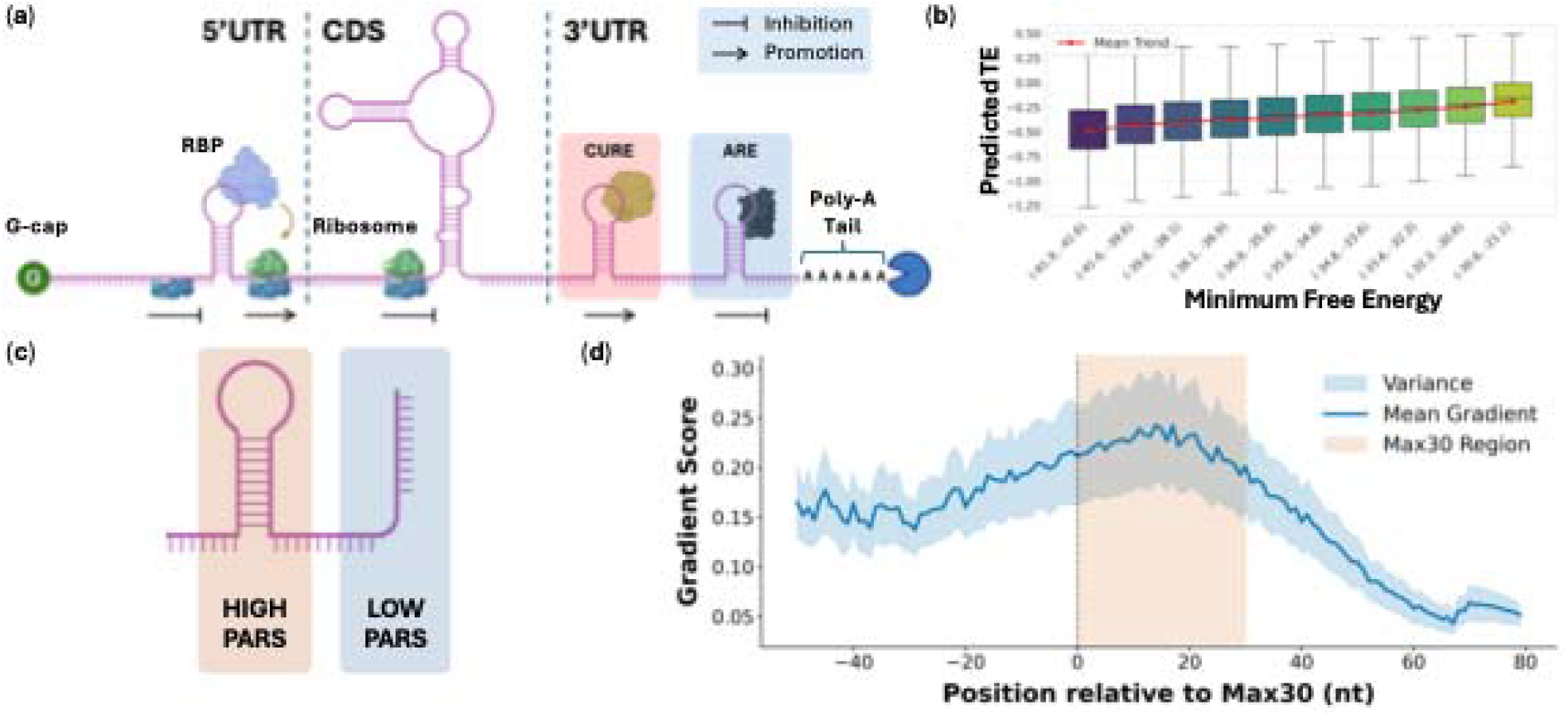
**(a)**Schematic of mRNA structural motifs and regulatory interactions. RBP refers to RNA-Binding Protein. **(b)**Distribution of TRIM-predicted TE across Minimum Free Energy (MFE) bins, with the median indicated by the black line. **(c)**High PARS scores (pink) indicate a higher likelihood of secondary structure (e.g., stem-loops) and Low PARS scores (blue) represent single-stranded regions with lower pairing probability. **(d)**Analysis of structural complexity and gradient significance. The distribution shows the mean gradient score across genome-wide analysis, and the start of the Max30 region is aligned to position 0.

From a thermodynamic standpoint, we calculated the Minimum Free Energy (MFE) for all sequences in the Cuperus dataset[5] using RNAfold and performed a correlation analysis (**Fig.3b**). Notably, in the low MFE region (<-40 kcal/mol), where structures are highly stable, the predicted TE is strongly repressed. The result suggests that TRIM has learned the biological rule that complex secondary structures inhibit translation.

We further incorporated PARS (Parallel Analysis of RNA Structure) to provide experimental ground truth[12,22]. PARS is a high-throughput enzymatic probing technique that quantifies structural complexity by distinguishing stem-looped regions from single-stranded RNA (**Fig.3c**).

To locate the focus of TRIM, we employed integrated gradients to determine the contribution of each base to the final TE prediction, defining this metric as the Gradient Score. Genome-wide mRNA sequences were aligned to the start position of a 30nt window with the highest PARS score (Max30 PARS Region). The average gradient score displays a distinct unimodal distribution, with the peak falling within the MAX30 region (**Fig.3d**). It indicates that TRIM automatically captures the regions of highest scanning resistance in the 5’UTR and adjusts its prediction accordingly.

### 3.4. TRIM deciphers regulatory patterns and interactions

The 5’UTR sequence, especially the region immediately upstream of the start codon, plays a decisive role in TE. To evaluate TRIM’s capability in deciphering combinatorial regulatory logic, we analyzed mutagenesis data of the RPL8A [7] (see Materials and Methods).

We first calculated the mean ΔTE across all sequences harboring a specific mutation at each position, and the data were grouped into high-TE (top 10%) and low-TE (bottom 10%) subsets to isolate robust regulatory motifs. The resulting heatmap of high-TE variants reveals that position −2 acts as a regulatory hotspot and the sequences with a Cytosine (−2C) consistently exhibit elevated TE (**Fig.4a**). Conversely, specific mutations in the low-TE subset tend to suppress expression (**Fig.4b**), demonstrating that TRIM effectively distinguished regulatory hotspots from the background context.

**Fig. 4:**
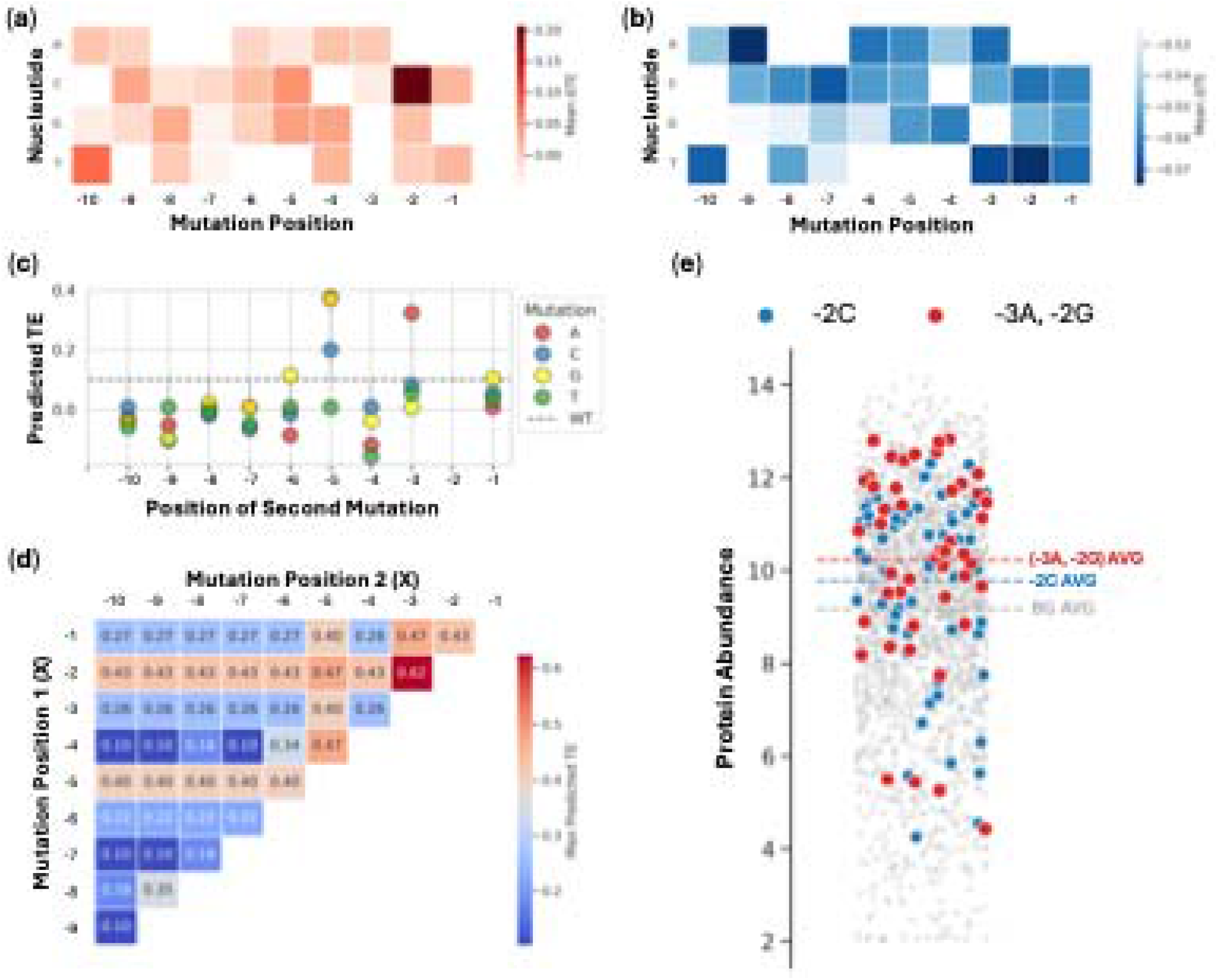
**(a**,**b)**Heatmaps of mean ΔTE across sequences harboring specific mutation within the (a)top10% (high-TE) and (b)bottom10% (low-TE) subsets. **(c)**Secondary mutation scanning on the (−2C) background. Each point represents the predicted TE of a sequence containing the (−2C) mutation plus one additional mutation. The dashed line represents the predicted TE of the sequence without mutation. **(d)**Heatmap of pairwise mutational scanning. Each block represents the maximum TE increase among all possible mutations for the corresponding position pairs. **(e)**Swarm plot of experimental protein abundance, showing the distribution of background controls(grey), single mutants (−2C, blue) and double mutants (−3A, −2G, red).

Identifying −2C as a primary driver, we conducted a secondary mutation scan (**Fig.4c**). Specific combinations, such as (−5A, −2C), (−5G, −2C), (−3A, −2C) exhibit synergistic effects that further promote TE, whereas pairs like (−4A, −2C) and (−4T, −2C) display antagonism.

We further expanded to a pairwise saturation scan within the [−10, −1]nt window (**Fig.4d**). Notably, while single mutations (−3A) and (−2G) yield lower predicted TE compared to (−2C), their combination (−3A, −2G) outperforms all dual mutations involving (−2C). This demonstrates that TRIM can identify non-additive effects, allowing the model to transcend the local optimum of single-point mutations and discover superior multi-site combinations.

Validation using experimental protein abundance supported our predictions (**Fig.4e**). To minimize noise, we excluded variants with high sequence deviation (number of extra mutant base >5nt were excluded). The mean protein abundance of sequences harboring the (−3A, −2G) double mutant significantly outperforms those with the single (−2C) mutation, and both exceeded the population average. These results are highly correlated with TRIM’s predictions, confirming its ability to decipher complex regulatory grammar.

## 4. Conclusion

In this study, we present TRIM, a transformer-based deep learning framework designed to accurately predict translation efficiency by integrating readily available transcriptomic and proteomic data. By synergistically modeling the 5’UTR, CDS and 3’UTR along with environmental features, TRIM achieves remarkable performance (Pearson *r* >0.9) without relying on costly and technically demanding ribosome profiling data. This significantly lowers the barrier for widespread application and allows for robust generalization across diverse datasets.

Beyond accurate prediction, TRIM demonstrates outstanding biological interpretability, effectively elucidating fundamental regulatory rules. The model precisely capturs the impact of mRNA secondary structure stability (MFE) and codon bias on translation dynamics. Notably, our interpretability analysis highlighted the non-linear interactions between regulatory motifs, such as the synergistic effects of upstream mutations (e.g., −3A. −2G) in the 5’UTR. Furthermore, *in silico* evolution experiments proved TRIM’s capability in protein engineering. The model not only optimized CDS sequence for enhanced expression but also spontaneously reproduced the *Translation Ramp* phenomenon, sacrificing local codon optimality near the start codon to ensure global elongation fluidity without any explicit supervision.

TRIM bridges the gap between sequence data and functional protein expression, serving not only as a predictive tool but also a generative engine for synthetic biology. Validation on the in vivo experiments has confirmed TRIM’s ability to detect and guide targeted mutations, suggesting the potential utility in upscaling the production of industrial enzymes or therapeutic proteins with minimal optimization cost. Furthermore, the bioinformatics analysis of UTR and CDS has also demonstrated that TRIM can effectively identify kinetic and thermodynamic regulatory patterns during various translation stages relying solely on sequence and abundance data. This proven ability might facilitate the translational regulation analysis in non-model species where functional genomics data are scarce.

Overall, TRIM offers a novel, interpretable deep learning approach that leverages attention mechanisms and biological prior knowledge to analyze the regulatory grammar of mRNA. Our pipeline yields reliable prediction and mechanistic insights using only sequence and available abundance data, providing a scalable solution for the broad field of computational biology.

## Acknowledge

This work was supported by the grant from the College of Biomedical Engineering, Fudan University.

